# Sexually dimorphic recombination can facilitate the establishment of sexually antagonistic polymorphisms in guppies

**DOI:** 10.1101/365114

**Authors:** Roberta Bergero, Jim Gardner, Beth Bader, Lengxob Yong, Deborah Charlesworth

## Abstract

Recombination suppression between sex chromosomes is often stated to evolve in response to polymorphisms for mutations that affect fitness of males and females in opposite directions (sexually antagonistic, or SA, mutations), but direct empirical support is lacking. The sex chromosomes of the fish *Poecilia reticulata* (the guppy) carry SA polymorphisms, making them excellent for testing this hypothesis for the evolution of sex linkage. We resequenced genomes of male and female guppies and, unexpectedly, found that variants on the sex chromosome indicate no extensive region with fully sex-linked genotypes, though many variants show strong evidence for partial sex linkage. We present genetic mapping results that help understand the evolution of the guppy sex chromosome pair. We find very different distributions of crossing over in the two sexes, with recombination events in male meiosis detected only at the tips of the chromosomes. The guppy may exemplify a route for sex chromosome evolution in which low recombination in males, likely evolved in a common ancestor, has facilitated the establishment of sexually antagonistic polymorphisms.

## Introduction

In a diversity of organisms, the sex chromosome pair have evolved suppressed recombination around the sex determining locus, sometimes extending across the entire chromosome ^1, 2, 3^ and recently reviewed in ^4, 5^. One prominent theory for the repeated, independent evolution of suppressed recombination is that sexually antagonistic (SA) factors at loci closely linked to a sex-determining locus establish polymorphisms, with one allele benefitting males and becoming associated with the male-determining allele at the sex-determining locus, while another allele that is favoured in females is associated with the alternative allele at the sex-determining locus ^6^. Such linkage disequilibrium (LD) between the alleles at the SA locus and the sex-determining region generates selection for closer linkage between the two loci ^3^. Although this model is plausible, and this process may be important in sex chromosome evolution ^7^, evidence supporting its operation is scarce. If recombination between the SA gene and the sex-determining region has become suppressed, so that the entire region co-segregates in genetic crosses, it becomes difficult to detect the existence of a separate gene with a SA polymorphism within the region.

The fish *Poecilia reticulata* (the Trinidadian guppy) is particularly suitable for studying whether SA polymorphisms indeed lead to recombination suppression, because SA selection has been shown to be ongoing in natural guppy populations ^8^. Male coloration factors are polymorphic within populations of guppies in many rivers in the Northern Range of mountains in Trinidad ^8^. Conspicuous male coloration is favoured by sexual selection, but also increases visibility to predators. Thus male coloration traits are sexually antagonistic: in males, the risk of predation may be outweighed by the mating advantage (particularly in up-river populations, where predation is less severe due to waterfalls preventing the main predatory fish moving up from down-river sites), whereas such traits give females no advantage.

The involvement of SA selection is inferred from genetic studies showing that the factors controlling male coloration elements are concentrated on the sex chromosome pair, which carries 79% of factors found in natural populations, as reviewed by ^8^, although only around 4% of the physical genome is represented by chromosome 12 (LG12) ^9^, which carries the sex-determining region ^10, 11^. Of the sex-linked male coloration factors, slightly more than half are fully sex-linked, while the others are partially sex-linked, being located at most about 10 centiMorgans (cM) from the sex-determining locus ^8^; these are expressed only in males ^8^. Together, these classical results suggest that SA polymorphisms may have arisen on this chromosome, and subsequently evolved either male-specific expression, or complete sex linkage (the two expected routes by which conflicts between the sexes can be resolved ^12^). Although SA selection clearly occurs in natural guppy populations, negative frequency dependent selection also contributes to maintaining SA polymorphisms, as rare male morphs have advantages in mating ^13^ and higher survival, ^14^.

A recent study in guppies ^15^ inferred that an extensive non-recombining region forms about half of the guppy sex chromosome pair, organised into two evolutionary strata similar to those described in other vertebrates, including mammals ^16, 17^, and birds ^18^. Specifically, independent evolution of recombination suppression in different upstream populations of guppies was inferred ^15^. Such strata are consistent with the SA polymorphism hypothesis for recombination suppression, but are not definitive evidence, because other evolutionary situations can potentially select for reduced recombination around sex-determining loci ^19^.

Here we describe results of genetic and population genomic approaches using fish from the same Trinidad guppy population as one studied by Wright et al. ^15^. In agreement with Wright et al., we detect regions showing linkage disequilibrium with the sex-determining locus, but we describe evidence that these do not represent evolutionary strata, but instead reflect LD with SA polymorphisms, as hypothesised in models of fully and partially sex-linked regions ^20^. We found at most only a few LG12 SNPs with fully sex-linked genotype configurations, with many variants apparently showing partial sex linkage, probably due to gene conversion.

Our genomic results strongly suggest that crossing over is very rare across most of the chromosome, with 97% of its gene content showing strong LD with the sex-determining locus. To further evaluate this interpretation, we carried out genetic mapping, which revealed sexually dimorphic overall recombination rates (heterochiasmy), with very different patterns of crossover localisation in the two sexes. In male meiosis, crossing over was detected only at the tips of all chromosome tested (chromosome 12 and three autosomes). The sex chromosome is therefore not unique in this respect, but, because it carries the male-determining locus, a large region that includes this locus will be transmitted exclusively through the male lineage, allowing a differentiated Y-specific region to evolve ^21^.

The prominent heterochiasmy found in a sexually dimorphic species is intriguing. Since the heterochiasmy is genome-wide in the guppy, it probably evolved before LG12 became the sex-determining chromosome, and not as a consequence of SA polymorphisms. Even so, it paved the way for an alternative, non canonical, evolution of the guppy sex chromosomes and for the enrichment of sexually antagonistic male colouration genes on the sex chromosomes.

## Results

### Population genomic tests for associations with the sex-determining locus

With the aim of defining the fully and partially sex-linked region of the guppy chromosome 12, we sequenced the complete genomes of 10 male and 6 female fish from a captive population of guppies derived from an Aripo river high-predation population (see Methods). The read mapping rate is slightly higher in females than males (97.638% versus 97.510%; Kruskal-Wallis statistic 6.80, P value = 0.009, Supplementary Table S1), suggesting that a small number of diverged Y-linked sequences might fail to be mapped.

We analysed F_ST_ values between the sexes for individual variable sites (see Methods) to test for associations between SNPs and the sex-determining locus ^20^. Considerably higher F_ST_ values were estimated for LG12 than for the other chromosomes (Figure 1), supporting its previous identification as the sex chromosome pair ^10^. As previously found ^15^, SNP density was also highest on LG12 (Supplementary Figure S1). However, the difference from other chromosomes was less clear than for F_ST_, not surprisingly, given that coding and non-coding sequences were not analysed separately, and diversity values will therefore depend strongly on the densities of coding sequences in different regions. Analysis of F_ST_ values does not suffer from this problem, because F_ST_ is a relative measure, estimating the proportion of variability that is between the two populations analysed; thus, both coding and non-coding SNPs can reveal associations with the sex-determining locus. Consistent with the F_ST_ analysis, Tajima’s D values for LG12 SNPs were higher in males than females (Supplementary Figure S2A).

**Figure 1.**
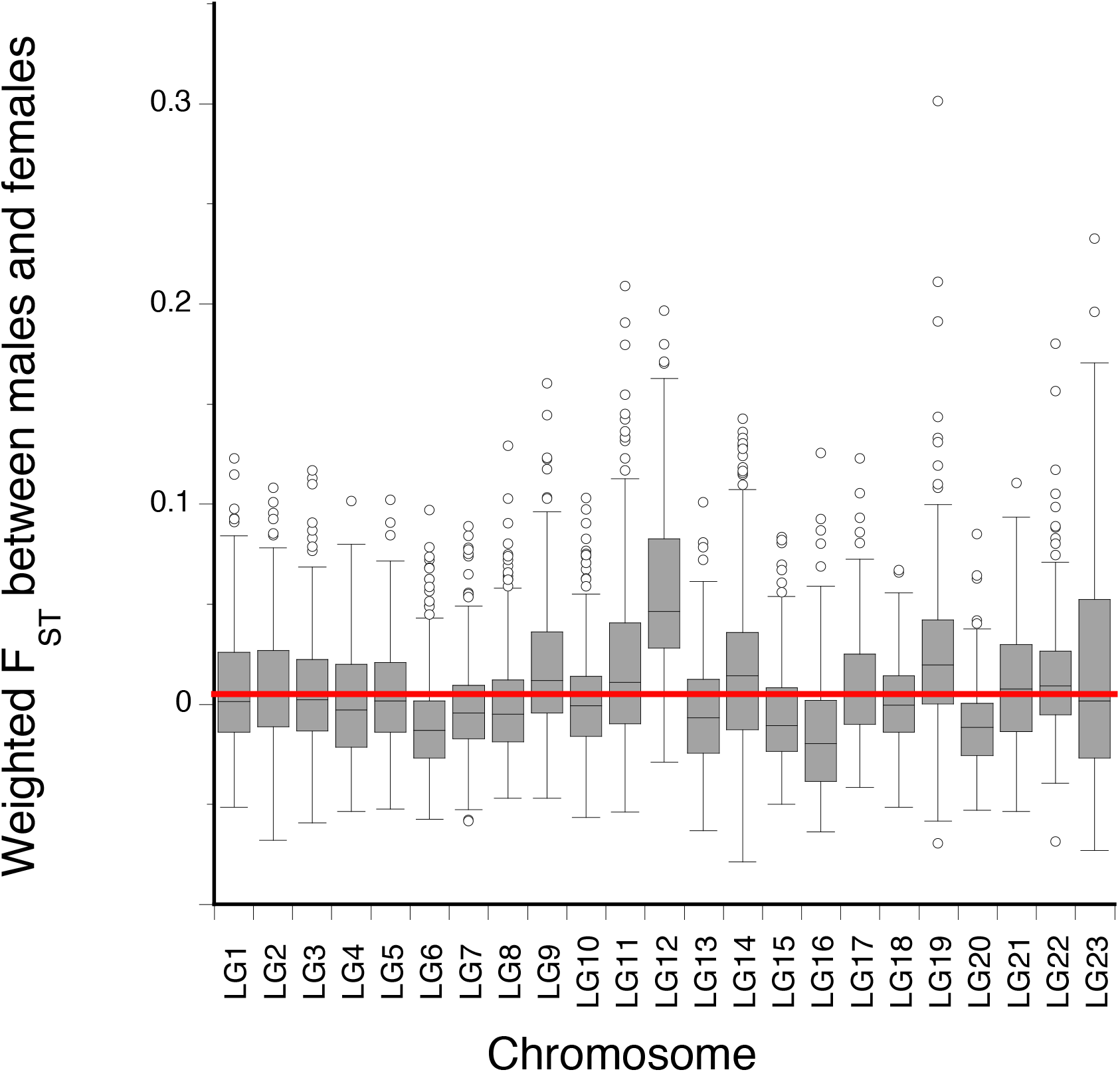
*F*_ST_ between males and females for each of the 23 guppy chromosomes. Only sites with genotypes for all individuals were included in the analysis. Each bar encloses 50% of the data, and lines within each box show the medians. The lines extending from the top and bottom of each box indicate ± 1.5 times the inter-quartile distance (and outliers are plotted as individual circles). The red horizontal line indicates the mean for all linkage groups other than LG12. We also estimated F_ST_ values for the autosomes, to test whether the differences observed could arise by chance. The value for LG12 differs highly significantly from the mean for each other chromosome (Kruskal-Wallace tests yield P values all < 0.001).

However, we did not detect any extensive fully sex-linked region on LG12. Instead, almost all variants on this chromosome appear partially sex linked. F_ST_ analysis of 50 kb windows across chromosome 12 (see Methods) revealed that no region had F_ST_ approaching the value 0.27 expected for fully sex-linked sites with our sample size. The first three megabases of the assembly had higher male-female differentiation than elsewhere on chromosome 12 (Figure 2A; Tajima’s D values were also higher in this region, see Supplementary Figure S2B), consistent with results from a captive population of fish from a Quare river source ^22^. However, this region also shows linkage disequilibrium (LD, estimated as *r*^2^ values, see Methods) in the pooled sample of fish of both sexes (Figure 2C), probably due to being close to the centromere (Figure 2D).

**Figure 2.**
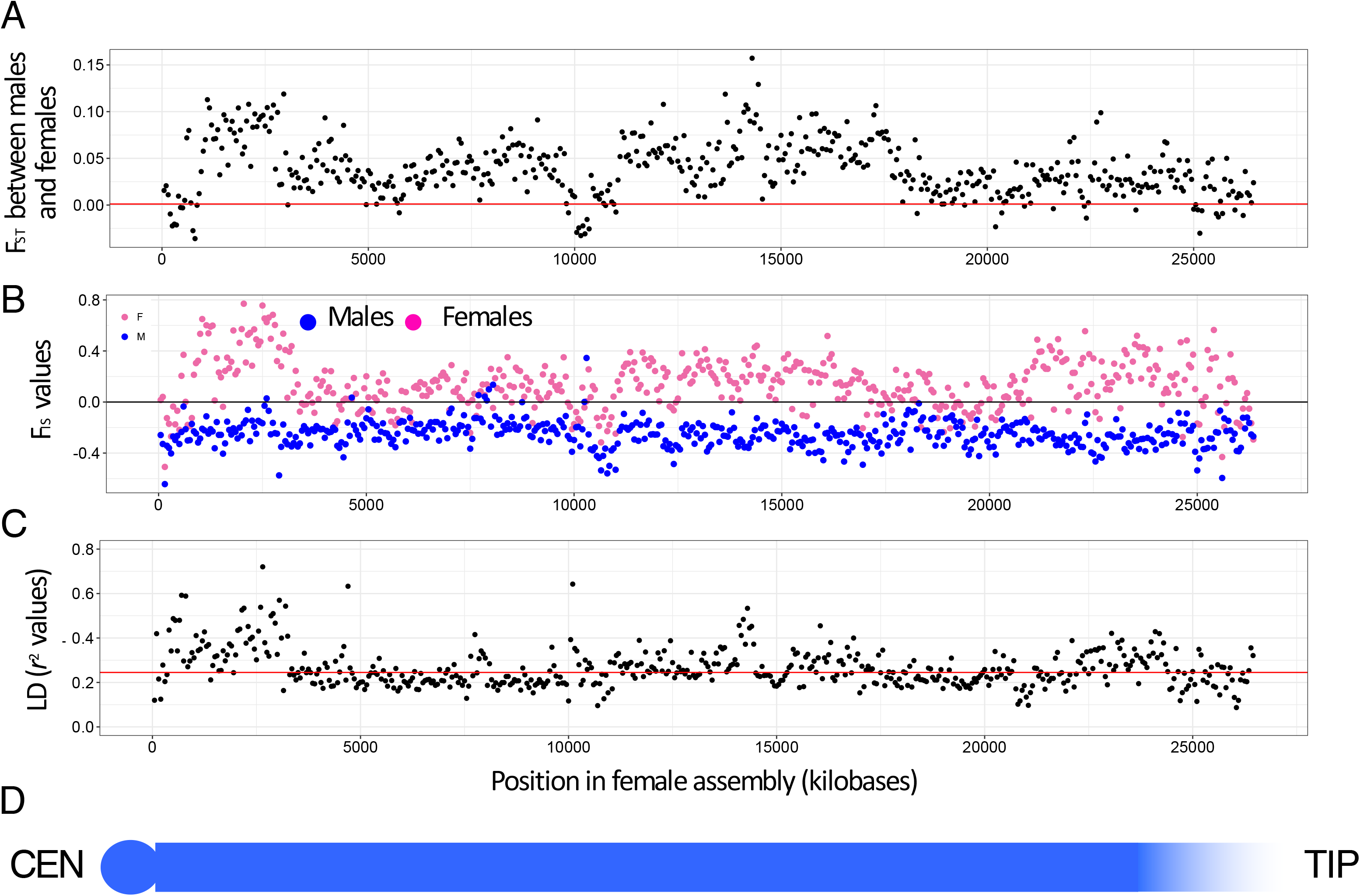
Evidence for sex linkage of SNPs on chromosome 12. **A**. F_ST_ values between males and females, based on 469,515 biallelic SNPs with genotypes for all individuals. The values plotted are mean values in 50 kb windows, and the horizontal line show the chromosomal mean values for F_ST_. Chromosome positions are based on the female genome assembly available in GenBank. **B**. F_IS_ values in males (blue) and females (red) separately, showing the more negative values in males in almost all 50 kb windows across LG12, with the possible exception of the tip near 26 Mb. The line indicates a value of F_IS_ of zero, as expected under Hardy-Weinberg equilibrium. **C**. Linkage disequilibrium (LD) between SNPs, estimated from diploid genotypes of both sexes using *r*^2^ values. To estimate the *r*^2^ values, SNPs were thinned to one SNP per 2kb, for sites with no missing data, and a minor allele frequency of 0.2. LD is high at the end of LG12 near 0 bp in the female assembly, supporting its assignment as the centromere end. Away from this end, *r*^2^ decays to a value close to 0.2. As the population studied here has a history of a small size at its founding (see Methods), and our sample size is small, LD is not expected to decay to very low values, even for independently segregating SNPs ^24^. The horizontal line represents the chromosomal mean values for LD. **D**. Schematic diagram of the chromosome, showing the centromere at the left (CEN), corresponding with the region of high LD, and a gradient of recombination rates in males at the tip (labelled TIP), where recombination occurs frequently.

Low diversity is a common cause of F_ST_ peaks ^23^. We therefore compared heterozygote frequencies in the two sexes by analysing F_IS_ values for individual SNPs (see Methods). In a fully sex-linked region, high frequencies of heterozygotes specifically in males are expected, with many sites expected to have F_IS_ = −1 in males, while females’ genotype frequencies should be close to the Hardy-Weinberg proportions (yielding F_IS_ values near zero, or positive values if the population is inbred). F_IS_ analysis indeed indicated sex linkage at many LG12 sites (Figure 2B), especially in the regions showing high male-female differentiation in Figure 2A. The peaks in F_ST_ values are therefore due to variants at intermediate frequencies that are commoner in males than females, supporting partial Y linkage.

Only 14 LG12 SNPs have genotype configurations compatible with complete sex linkage (labelled “XY SNPs” in Figure 3). Among these, ten clustered in a 15.6 kb region between positions 2,937,371 and 2,952,959 bp in the female assembly surprisingly, given the expectation that the sex-determining region is near the tip of LG12 ^10, 11^. Three other such SNPs were in a much larger region at a considerable physical distance from this cluster (> 574 kb between positions 15,268,306 and 15,843,036), and one was located at position 7,677,458. Variants at many sites are shared between the two sexes, and biallelic sites with male homozygotes for both alleles were abundant, which cannot occur without recombination. No SNP with the genotype configuration expected under full sex linkage was found within the 3 Mb region identified as the older stratum and inferred to be found in all guppy populations ^15^. Our sample size does not permit us to conclude firmly that any SNP is fully sex-linked, and therefore our analysis probably over-estimates the number of fully sex-linked SNPs on LG12.

**Figure 3.**
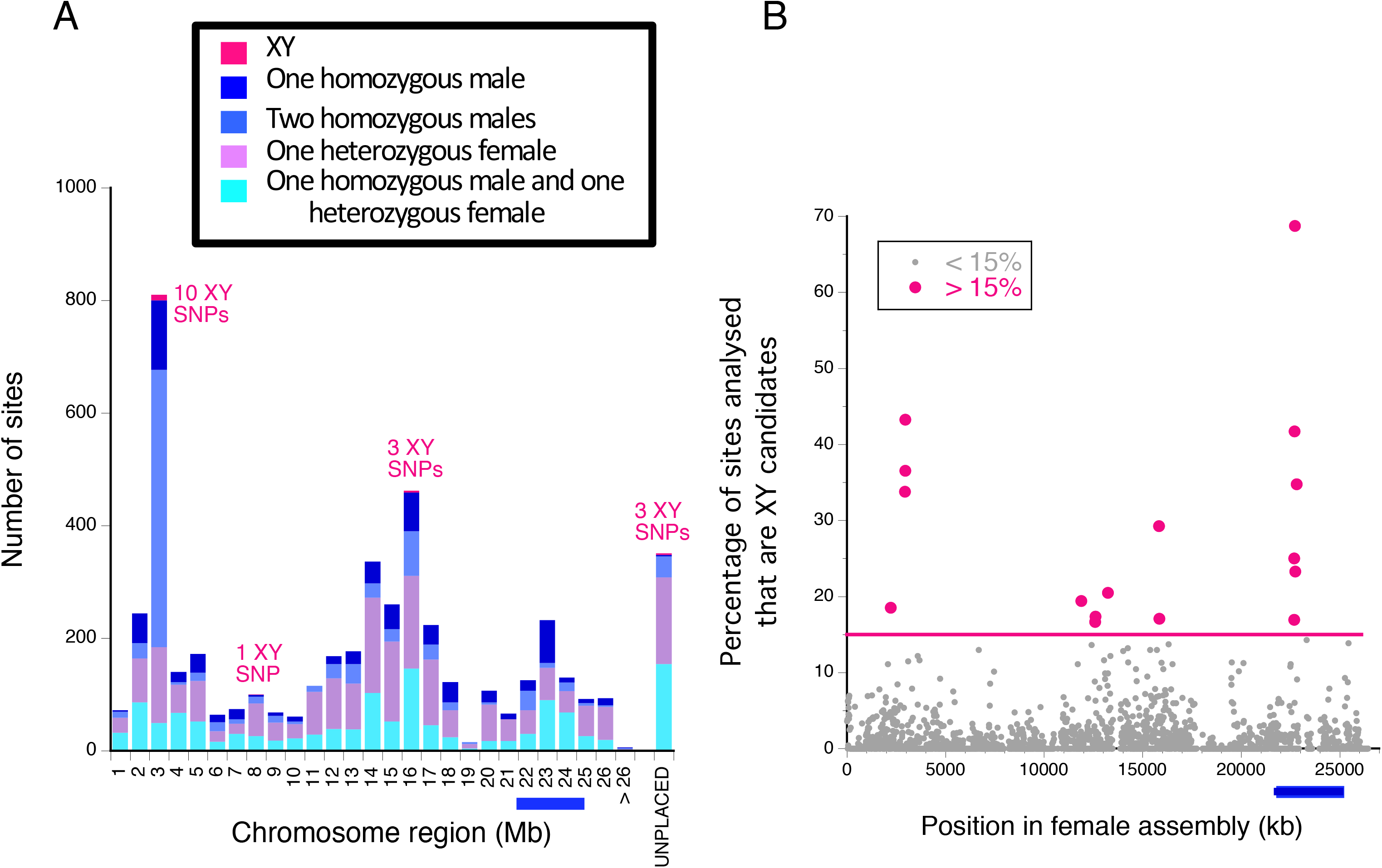
Numbers and proportions of of SNPs on the *P. reticulata* chromosome 12 showing different levels of evidence for sex linkage, based on the genotype configurations in the two sexes. The blue bars under the x axis at the right-hand side indicate the region previously suggested to be an old evolutionary stratum ^15^. A total of 344,848 non-singleton variable sites from chromosome 12, and 492 unplaced scaffolds with at least 100 variable sites, were analysed. Sites in the 5 categories defined in the Methods section as most strongly suggesting sex linkage are concentrated in three regions of the chromosome. These scattered footprints of sex-linkage are not due to errors in the assembly, as our female genetic map of LG12 (Figure 3 below) indicates the same order as in the female reference genome sequence, with one small exception near 22 Mb. **A**. The locations (in megabase regions) of sites with genotype patterns expected under complete sex-linkage, and sites with slightly deviating configurations (see Methods). **B**. Proportions of sites in 10 kb windows that fall into any of the categories shown in part A. The line separates the small number of windows in which more than 15% of sites analysed have any of the genotype configurations shown in part A, which are also shown in red, while the proportions in the other windows are shown as grey dots.

We also counted the number of sites with signals of complete or partial sex linkage found on the autosomes, to estimate the number expected by chance. Consistent with our observations for the F_ST_ values, the other chromosomes show markedly smaller numbers of sites with signals of complete or partial sex linkage; in total, only 21 SNPs on the 22 chromosomes other than LG 12 had genotype configurations expected under complete sex linkage (heterozygous in all 10 males, and homozygous in all 6 females studied), and only 3 autosomes had more than two such SNPs; these are probably mostly false positives. The over-representation of LG12 is therefore striking; the random expectation, based on either the number of SNPs analysed, or the estimated chromosome sizes, is that LG12 should carry about 4% of such SNPs, while the proportion we observed is 40% (in 10,000 trials of 35 sites assigned randomly to 23 chromosomes, the number found on a single chromosome never exceeded 7). Moreover, only three further SNPs with this pattern were detected among the unplaced scaffolds (Figure 3).

Detailed analysis of the genotypes of SNPs on the guppy sex chromosome pair revealed numerous sites with only one or two genotypes deviating from complete sex linkage (Figure 3A). These are particularly abundant in the three regions that also have the apparently fully sex-linked SNPs (Figure 3A), and form high proportions of the SNPs in 10-kb windows (Figure 3B) in the same three regions of the assembly. These footprints of sex-linkage are therefore not produced by regions with unusually high SNP densities. However, although one of these regions lies between 22 and 25 Mb, where an old stratum has been inferred ^15^ (Figure 3B), not a single SNP with the genotype configuration indicating complete sex linkage was detected in the region. Overall, these results suggest that there is a very small region showing full sex-linkage, perhaps like the single gene sex-determining systems known in some other fish ^24, 25, 26^, and a recent new analysis in the guppy also infers a small ancestral fully sex-linked region, using the same fish previously used to infer the 3 Mb old stratum ^27^. An analysis to test for male hemizygosity also found no evidence that this is common on LG12 (Supplementary Figure S3A). Nor did we detect coverage differences between the sexes, or evidence for male hemizygosity, in the region where the previous analysis ^15^ detected lower coverage in males (Supplementary Figures S3B and 4), although our coverage analysis differs from theirs (see the Methods section). However, the region just proximal to 3 Mb, with multiple sites heterozygous for male-specific variants, shows higher coverage than the flanking regions, indicating that the male-specific SNPs are in a duplicated copy of this non-coding region.

Despite the fact that genotype configurations expected under complete sex linkage were infrequent overall, a major sub-set of the individuals studied had genotypes that conform with this configuration at many sites. Deviations from this configuration were largely due to three individuals, two males that were homozygous for the commonest variant in females at many LG12 sites, instead of being heterozygous, and one female that often had the heterozygous genotype found in males (Supplementary Figure S5). The failure of our analysis to detect regions with genotype configurations compatible with complete sex linkage is therefore not due to genotyping errors, because this should affect all individuals roughly equally. When these three individuals are excluded, the F_ST_ values increase, and the remaining Y haplotypes include many sites with configurations compatible with complete sex linkage (although the reduced sample size means that this is not definitive evidence for any region being completely sex linked). F_ST_ values still approach the value expected for completely sex-linked SNPs only in two of the regions showing the footprints of sex-linkage mentioned above (red boxes in Supplementary Figure S6), but again not in the region of the inferred old stratum ^15^.

These genotypes are recombinants between the commonest X and Y haplotypes, though they were not produced by recent recombination during captivity, as the sites with genotypes incompatible with complete sex linkage are found throughout the entire LG12 assembly, rather than forming extended haplotype regions separated by evident recombination events. To investigate these individuals further, we applied principal components analysis (PCA) to our SNP data. As shown in Figure 4A, two individuals that are outliers for LG12 SNPs also cluster together for SNPs on LG1 and LG9, while one does not. These results suggest migration from a closely related population, most likely an Aripo up-river site. Such migration has been inferred in other studies ^28, 29^. For LG12, the first two principal components explain only 17 and 13% of the variation, and somewhat smaller values for the two autosomes, consistent with the migrants being from a closely related source population.

**Figure 4.**
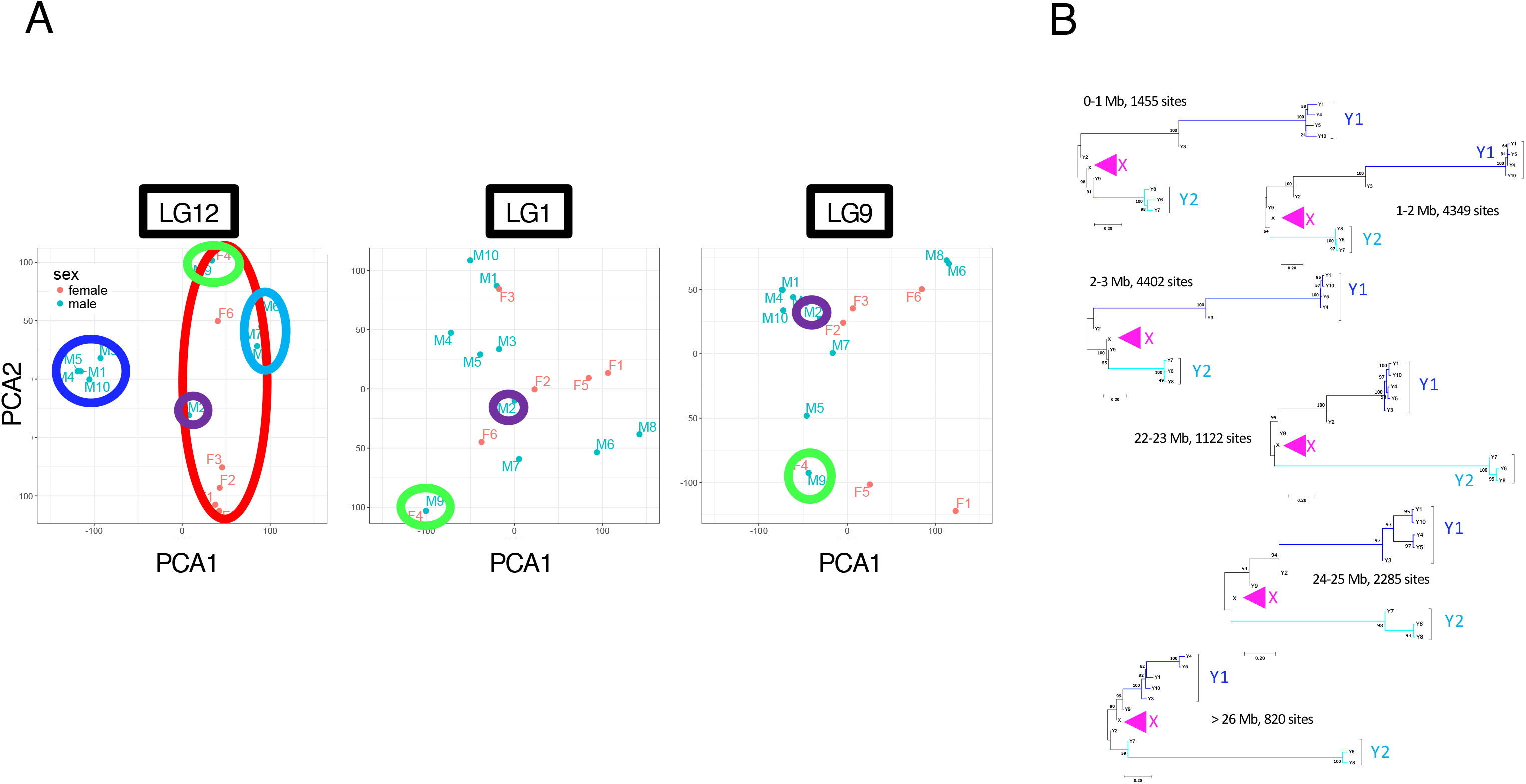
Principal components analysis of SNPs that were genotyped in all 16 individuals in our sample, for LG12 and two autosomes. In the PCA for LG12, the red outline encloses the females, whereas the two areas outlined in different shades of blue are males with distinct Y haplotypes; purple and green outlines enclose the three individuals whose genotypes for LG12 SNPs deviate from those expected under complete sex linkage (see Figure S5). Green indicates male 9 and female 4 in Figure S5, and purple indicates male 2. In the PCA for LG1 and LG9, the two individuals (male 9 and female 4) clustered together in the PCA for LG12 also cluster together for these two chromosomes, and appear as outliers from the other individuals, while male 2 does not cluster with these two individuals. **B**. Neighbour-joining trees for different regions spanning LG12, based on biallelic sites with homozygosity in all 6 females sequenced, but with heterozygosity in at least 2 males. The trees are rooted with the inferred X sequence at these sites, based on the female states. The numbers on the branches are bootstrap values based on 500 replications.

**Figure 5.**
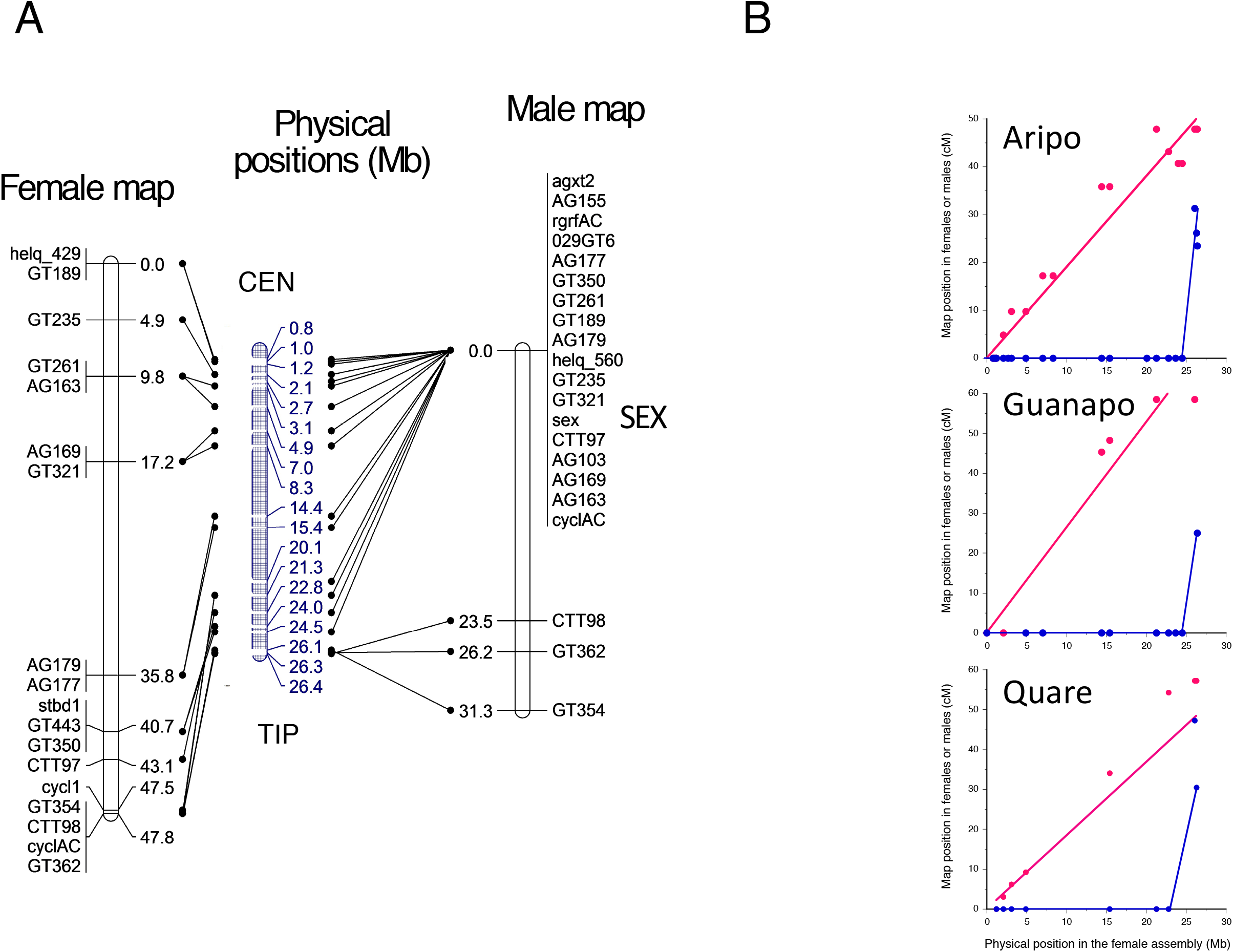
Genetic mapping results for chromosome 12 based on full sib family from the Aripo laboratory population and natural populations from the Guanapo and Quare rivers (see Methods). **A**. Genetic maps for male (right-hand side), and female meiosis meiosis (left); the physical positions of the markers in the female assembly (estimated LG12 total physical size = 26.5 Mb ^9^) are shown in the centre. The *cyc1* gene was mapped previously ^30^, using a marker named 0229, and is the closest marker to the SEX locus in their map (which was merged for the two sexes). In situ hybridisation of a BAC clone, 34-K02, which carries this gene, located it near one tip of the *P. reticulata* sex chromosome pair ^10^, consistent with the location in our female genetic map. In our male map, this gene (marker *cyclAC*) co-segregates with markers that did not recombine with the sex-determining locus (indicated by SEX in the figure). These markers span the physical region that includes 10 SNPs with genotype configurations expected for complete XY linkage. **B**. Marey maps showing the consistent marked difference between male (blue) and female meiosis (red) in the Aripo family family shown in A, together with results from two families from parents sampled in natural down-river populations (see Methods). The genetic map locations estimated in female meiosis in all three families increase almost linearly with the physical positions in the assembly (red lines); in the Aripo family, R^2^ = 0.95, in the Guanapo family R^2^ = 0.93, in the Quare one R^2^ = 0.65. The difference between the female and male recombination rates is highly statistically significant in several independent intervals of LG12 in the Aripo river family, and in the single interval where this comparison is possible in the Guanapo family (Supplementary Table S4).

The PCA analysis (Figure 4A) also suggests the presence of two different Y haplotypes, which could also contribute to the low intersexual F_ST_ values we observe across LG12, along with rare recombination, which (as explained above) is indicated by the presence of many biallelic sites with male homozygotes for both alleles. To further test for the existence of two haplotypes, we examined all biallelic sites that were homozygous in all six females in our sample, and recorded how many males were heterozygous for another allele. Many more sites had 3 or 5 males heterozygous than any other configurations, supporting the presence of two groups of males with differing Y haplotypes among the eight males other than those with the likely immigrant and recombinant Y chromosome genotypes discussed above. We used these sites where the phase of variants can be inferred to make trees; these again show that these haplotypes extend across all of the chromosome, with strong bootstrap support, except perhaps for the tip region (Figure 4D). They also suggest that the two Y haplotypes formed soon after the X haplotype was established, which indicates the time when LG12 became a sex-determining chromosome.

### Genetic mapping

Previous high density genetic mapping of guppies indicated a linkage group carrying the sex-determining locus ^30, 31^. This corresponds to chromosome 12 in the genome assembly ^9^. Cytogenetic studies in domesticated guppies ^11^ revealed that crossovers in male meiosis localise near the chromosome tips, consistent with crossover locations detected by genetic mapping ^30^. For the XY pair, most cytologically detected crossovers were in the terminal 15% of the chromosome (most distal from the centromere), corresponding to about 4 megabases of the female assembly. A previous sparse genetic map using AFLP markers ^32^ did not detect crossover localisation in male meiosis.

We estimated separate genetic maps in male and female meiosis, and compared their crossover patterns, in full-sib F1 families from individuals from within-population crosses from the same Aripo river population as used for the population genomic analyses described above, and from wild-caught fish from the Guanspo and Quare rivers (see Methods), and mapped microsatellite and SNP markers, many of them informative for both male and female meiosis (Supplementary Table S2). In male meiosis of our families, most chromosome 12 markers co-segregate with the sex phenotype, confirming that chromosome 12 carries the sex-determining locus in all our male parent fish (Figure 4). Cosegregation of markers was observed even for the centromere-proximal region, consistent with having found male-specific variants in this region from our genomic data, and implying that recombination between this region and the sex-determining locus is infrequent.

The male and female meiotic maps differ in both the recombination rates and patterns. In male meiosis, recombination with the sex -determining locus was detected only for markers located in the terminal megabase of LG12, suggesting that the recombining tip region may be even more restricted than estimated cytogenetically ^11^. In contrast, recombinants were detected throughout chromosome 12 in female meiosis (Figure 4A and B), and the order of the markers in the female genetic map agrees well with their order in the female assembly ^9^, confirming its reliability, and showing that the markers are all LG12 sequences. In the Aripo family, the numbers of recombinants differ significantly between the sexes in three independent intervals, and in males the terminal interval differs highly significantly from that in the rest of LG12 (Supplementary Table S4). In the Guanapo and Quare families the differences are also significant.

To test whether heterochiasmy is restricted to LG12 (and might have evolved because this is the sex chromosome) or is a genome-wide feature of this species, we also mapped three autosomes (1, 9 and 18). Again, recombination events were detected in most intervals tested in female meiosis (Supplementary Figure S7), whereas, in male meiosis, crossovers were confined to the two most distal megabases of each chromosome.

## Discussion

The guppy sex chromosome system raises two interesting questions: how just 4% of the genome can carry so many male coloration genes (and possibly genes for other sexually antagonistic traits), and whether SA polymorphisms in the partially sex-linked region(s) are selecting against recombination with the sex-determining locus. Our results provide an answer to the first question. With a sexually dimorphic crossover pattern such as we observe, a chromosome that acquires a male-determining gene will immediately behave genetically as a Y chromosome without closer linkage evolving under selection against recombinant genotypes ^21^. The recombination pattern that we find genome-wide in male guppies will therefore have resulted in almost the entire chromosome experiencing sex-linkage, as soon as a male-determining factor appeared on it, facilitating the establishment of SA polymorphisms. At loci with such SA polymorphisms, male-benefit alleles are predicted to be found in association with the male-determining allele at the sex-determining locus, leading not only to linkage disequilibrium (LD) between alleles at the two loci, but also to LD between neutral variants located close to the SA locus and the sex-determining locus. As such associations depend on close linkage ^20^, they could occur across much larger genome regions in a species such as the guppy, where male crossing overs are restricted to small chromosome regions. The whole chromosome will be passed from fathers only to their sons, so that new mutations that arise on that chromosome will exhibit Y-linkage. If recombination is very rare between an XY chromosome pair, SA polymorphisms could become established anywhere in the partially (but nearly completely) sex-linked region, even at loci physically distant from the sex-determining locus. This can account for the inferred enrichment for male coloration genes (see above), and for genes likely to have functions in pigmentation processes ^27^. Such polymorphisms are expected to produce footprints of ongoing balancing selection on the sex chromosomes ^20^, including the observed elevated F_ST_ values between the sexes, when compared with other chromosomes.

The LD we detect on this chromosome pair, and the evidence for two distinct Y haplotypes in our sample (Figure 4), may therefore represent the expected footprints of sexually antagonistic polymorphisms, given that guppy male coloration factors are concentrated on this chromosome pair ^10^. Moreover, the observation of distinct Y haplotypes strongly suggests the action of frequency-dependent selection, as such polymorphisms are unlikely to be maintained in its absence ^33^. If SA selection is indeed involved, crossover localisation might evolve to reduce recombination.

The rarity of recombination across most of the XY sex chromosome pair probably largely reflects an ancestral state with chiasma localisation to the chromosome tips in males. The finding of sexual dimorphism of recombination rates and patterns in guppies and other fish species ^30, 31, 34^, as well as in amphibians ^21^ suggests that such differences are widespread in lower vertebrates. Several evolutionary reasons for heterochiasmy have indeed been proposed and modelled ^35^.

The prominent heterochiasmy found in guppy is also similar to the crossing-over patterns in *Rana temporaria* ^36, 37, 38^. Occasional sex-reversals in these frogs create XY females, allowing rare recombination events between the X and Y chromosomes ^39^, probably accounting for breakdown of associations between the male-determining locus and genetic variants in some *R. temporaria* populations, whereas other populations show male-specific haplotypes.

We now turn to the second question, namely whether SA polymorphisms have actually resulted in the evolution of less crossing over than occurred before these polymorphisms were established on the guppy sex chromosome. A recent paper concluded that recent recombination suppression events have occurred in low-predation populations, where selection against expression of male coloration traits by females is weaker. The authors concluded that the guppy Y chromosome carries an ancestral fully non-recombining region as large as 3 Mb, and a second, younger stratum evolved in up-river populations, forming a 10 Mb fully sex-linked region of this 26.5 Mb chromosome ^15^. According to this interpretation, our high-predation population should lack the younger stratum and should only have the inferred 3 Mb old non-recombining stratum surrounded by a terminal ∼1.5 Mb PAR from one side, plus a centromere-proximal recombining PAR of around 20 Mb from the other side. On the contrary, no crossover was detected for this centromere-proximal region in male meiosis of our three mapping families, thus clearly indicating that recombination must be rare in this region. More importantly, our population genomic results also show that many sites across the entire chromosome show LD with the sex-determining locus, which implies that recombination has been very rare over the past history of the individuals sampled.

It remains to understand what produces rare recombination in the guppy sex chromosome. Sex reversals appear to be very rare in guppies ^40^, but could allow rare XY reciprocal recombination in guppies, as in *R. temporaria* ^39^. Alternatively, the rare recombination we detect may represent gene conversion. In several organisms, gene conversion occurs in regions that do not undergo crossing over ^41^, and could occur in the guppy chromosomes, including the X and Y pair, during male meiosis ^12^. This can reconcile the overall picture of strong associations with the sex-determining locus, and distinct Y chromosome haplotypes, with clear evidence that recombination occurs.

In conclusion, if SA polymorphisms have selected for close linkage with the guppy male-determining gene, and led to lower recombination rates in low predation populations, as suggested ^15^, the mechanism is likely to have involved changes in crossover localisation; discrete recombination suppression events, such as inversions, creating evolutionary strata seem unlikely. Differences in crossover localisation are suggested by the observation that univalents were seen only for the XY pair, suggesting that this pair have more terminal localization of chiasmata than the autosomes ^42^. Larger numbers of meiotic products should be studied by genetic mapping in the future, and more metaphases should be examined for presence of univalents, to test whether chromosome 12 differs from the other chromosomes in the pattern or degree of localization of crossing over in male meiosis. Families from low-predation populations should also be studied to test whether crossover patterns in male meiosis differ genetically between populations, and whether any differences are specific to chromosome 12. Also, population genomic data from larger samples can test the surprising previous interpretation ^15^ of lower reciprocal recombination rates in low predation populations. If LD on chromosome 12 breaks down nearer to the centromere than in our high-predation sample, it will imply the opposite, namely that low predation populations have higher reciprocal recombination rates, consistent with the genetic data so far available ^43^.

## Methods

### Fish samples

Our study used a captive population derived from a sample collected by D.P. Croft (University of Exeter) in the lower part of the Aripo river in Trinidad (high predation zone, N 10°. 39.031; W 61°13.404) in March 2008. Approximately 200 fish were used to found the population (100 males and 100 females). The fish have been maintained in a large population at Exeter, in 6 pools (3 x 1.5 x 0.6m). The pools house over 12,000 fish, and individuals have been regularly moved between pools, to maintain the absolute population size as large as possible in stock ponds. The fish used for sequencing and genetic mapping came from a colony founded at the Cornwall campus in February/March 2013 from 600 adults from this large population, and maintained with a population size of at least 1,000. To maintain a high population size, the population was split across multiple tanks, with fish regularly exchanged between tanks, and without imposing any selection or inbreeding (other than the naturally occurring selection, including through female choice). Our sample was chosen to provide sequences from 22 X chromosomes and 10 Y chromosome. Fish were humanely euthanised by terminal anaesthetic in a solution of Tricaine methanesulfonate (MS222) buffered to neutral pH with sodium bicarbonate, and preserved in EtOH at −20 C.

#### Genome sequencing and analyses

Genomic DNA was extracted from 10-12 mg of caudal tissue from 10 males and 6 females, using the DNeasy blood&tissue kit (Qiagen, Hilden, Germany) and RNase-treated according to the manufacturer’s recommendations. Paired-end sequencing libraries were constructed from 3-5 µg of RNase-treated DNA and sequenced as 150-bp paired-end reads, and complete genome sequences were obtained from all 16 individuals (Supplementary Table S1 provides details of the insert sizes and other statistics).

For each fastq file, reads were trimmed to remove adapters and primers, along with poor quality bases, using cutadapt (version 1.8.3), using parameters -m 35 -q 301 ^44^. The resulting filtered sequences were mapped to a publicly available reference female *Poecilia reticulata* genome sequence assembly (available at GenBank under accession number GCF 000633615.1). Reads were aligned against the reference genome using bwa mem (version 0.7.13) ^45^ with parameter -M which marks split alignments as secondary so that they can later be excluded by downstream tools. Duplicates were marked using Picard tools (version 2.8.1) (http://broadinstitute.github.io/picard) and excluded. The minimum mapping rate among the 16 guppy samples was 97.41%, and the coverage, estimated for all sites using SAMTools (v 0.1.19), was rarely below 50 (Supplementary Table S3); note that this coverage analysis differs from that of ^15^, who excluded male reads not perfectly matching the female assembly, in order to use lower male coverage to detect diverged Y-linked sequences). Moreover, the mapping rates were similar for males and females (Supplementary Table S1).

SNPs in the resulting files in BAM format were called using the GATK pipeline (version 3.7.0). SNPs and INDELS from each sample were called using HaplotypeCaller with the following parameters: emitRefConfidence GVCF, –genotyping mode DISCOVERY, -stand call conf 30, –variant index type LINEAR, –variant index parameter 128000. Further joint genotyping was performed using GenotypeGVCFs with the default parameters, followed by use of SelectVariants ^46, 47, 48^ to select all variants with minimum depth of coverage (DP) of 20 and quality (QUAL) 30. VCF tools (version 0.1.15) ^48^ was used to generate separate VCF files for INDELS and SNPS from the initial vcf files, using parameters –keep-only-indels and – keep-indels respectively. SAMtools (v 0.1.19) was used to estimate read depths of individual sites in the assembly, for each fish, and the resulting values were analysed using Python scripts. Coverage was high for the vast majority of sites, including those with the SNPs analysed, in all individuals (Supplementary Table S3 and Figure S**4**).

Population genomic analyses of SNPs, including estimates of F_ST_, Tajima’s D, and linkage disequilibrium (LD, which was quantified as *r*^2^, using analyses of the diploid data from both sexes generated by sequencing, without inferring phase) were done in VCF tools ^48^, as were F_IS_ estimates. The F_ST_ estimates were for individual variable sites, using the approach of ^49^. The results were plotted in non-overlapping windows of 50 kb. For a finer scale view of heterozygote frequencies, F_IS_ values were also estimated for each biallelic site, using a Python script. Principal Components Analysis (PCA) was done using the Tassel software ^50^.

To determine the extent of any regions exhibiting complete sex linkage, the genotype configurations of all variable sites with complete genotype information in all individuals, and no more than two alternative bases (biallelic SNPs), were examined. Such analysis allows us to identify sites with genotypes expected under complete sex linkage, i.e. sites with an allele fixed in the male population and absent in the female population, as well as sites showing partial sex linkage. Sites indicative of partial sex linkage were classified in the following arbitrary categories, and shown in Figure 3a: (i) at most 2 males homozygous for the inferred X-linked variant, (ii) at most one female heterozygous for the inferred Y-linked variant, (iii) at most one male homozygous for the X variant and one female heterozygous for the Y variant. As our sample includes sequences from 22 X chromosomes, X-linkage can be readily inferred. The alternative allele was then inferred to be Y-linked if present in most or all of the males. Finally, to test whether these partially or fully sex linked sites suggest clustering in certain individuals (Supplementary Figure S5), we used a single inferred X haplotype from each female individual, and a Y haplotype from each male; the tree is based on nucleotide p-distances analysed in MEGA software version 7 ^51^. The same procedure was used for the clustering shown in Figure 4D, based on sites with no variation in the females in our sample.

#### Genetic mapping

A full-sib family of 42 individuals (25 males and 17 females) was made using parents from the same population used for the genome resequencing, using as markers several microsatellites and SNPs in genes (Supplementary Table S2). For chromosome 12, the markers spanned most of the assembled chromosome, between 1 Mb and 25.6 Mb, with an average density of one marker per 1.1 Mb in the Aripo family. Microsatellites were genotyped by capillary electrophoresis using an ABI 3730 Sequencer (Applied Biosystems). SNPs from gene markers were either genotyped using RFLP assays or by direct Sanger sequencing. Genetic distances were inferred using JoinMap v. 4.0 ^52^, with a minimum LOD score of 3, using a regression mapping procedure, the Kosambi mapping function and a goodness-of-fit jump threshold of 5.0. In male meiosis, recombination frequencies between markers at the chromosome tips and the sets of co-segregating markers elsewhere in the physical assemblies of the chromosomes exceeded 30 cM, producing LOD scores below the threshold of 3. Linkage of these markers was supported by linkage inferred in female meiosis.

Genetic mapping was also done for two further families, made from parents collected in natural high-predation populations in the Guanapo and Quare rivers (GPS coordinates 10° 39' 28" N, 61° 15' 13", and 10° 39' 53" N, 61° 11' 35", respectively); it has been proposed that the Quare population may represent a different species, *P. obscura* ^53^, but other studies have not found such populations to be differentiated ^28^. The Guanapo family included 18 male and 11 female progeny, and the Quare family has 20 males and 15 females.

We also compared numbers of recombinants in meiosis of the male and female parents of all the families, using 2-tailed 2 x 2 contingency table analyses to test whether the sexes differ in their recombination. To make paired comparisons, we used intervals in which the physical distances were associated with multiple crossovers in female meiosis, and where both the markers defining the interval were genotyped in both parents of the cross (Supplementary Table 4).

#### Data availability

The sequence BAM files are available in the European Nucleotide Archive (STUDY_ID PRJEB22221).

## Acknowledgements

The project was supported by ERC grant number 695225 (GUPPYSEX). Genome resequencing and bioinformatics analyses (sequence alignments and production of BAM and VCF files) were carried out by Edinburgh Genomics, The University of Edinburgh. Edinburgh Genomics is partly supported through core grants from NERC (R8/H10/56), MRC (MR/K001744/1) and BBSRC (BB/J004243/1). We thank the University of Exeter animal technicians for their support with fish husbandry, and colleagues Konrad Lohse, Darren Obbard, and Alex Twyford at Edinburgh, and Alastair Wilson at Exeter, for discussions and comments on the manuscript.

## Author information

### Affiliations

Institute of Evolutionary Biology, School of Biological Sciences, University of Edinburgh, Charlotte Auerbach Road, EH9 3LF, U.K.

Roberta Bergero, Jim Gardner, Beth Bader, Deborah Charlesworth

Centre for Ecology and Conservation, University of Exeter, Cornwall, TR10 9FE, UK.

Lengxob Yong

## Contributions

DC designed the study with input from RB. LY produced the guppy genetic cross and families. JG and BB performs all molecular biology experiments, including mapping genotyping. RB analysed the data with input from DC. RB and DC supervised all aspects of the study. RB and DC collaborated to write the paper.

**Corresponding authors**: Correspondence to Roberta Bergero or Deborah Charlesworth

